# Systems serology of responses against tumor antigens in ovarian cancer reveal disrupted Fc-mediated immunity

**DOI:** 10.64898/2026.04.25.720834

**Authors:** Michelle Loui, Meera Trisal, Laura B. James-Allan, Scott D. Taylor, Het Desai, Gabriella A. DiBernardo, Allison Brookhart, Yun-Rong Ting, Jessica Gebraeel, Neda Moatamed, Pamela K. Kreeger, Sanaz Memarzadeh, Aaron S. Meyer

## Abstract

High-grade serous ovarian cancer (HGSOC) represents 75% of ovarian cancer cases and 80% of deaths, with most patients relapsing despite initial treatment response. The limited effectiveness of immunotherapies in HGSOC indicates urgent need for novel therapeutic approaches. HGSOC patients produce tumor-binding autoantibodies (TBAs) with high tumor selectivity. Since effective antibody-mediated tumor cell killing requires Fc domain interactions with immune cells, we hypothesized that, although TBAs recognize tumor cells, they might still poorly elicit cell killing responses. Using a systems serology approach, we profiled TBA subclass and biophysical interactions with Fc receptors in HGSOC, comparing them to antiviral antibody responses. TBAs were consistently identified within ascites and serum and were heterogeneous in subclass composition. However, TBAs consistently lacked the capacity to bind FcγRIIIa despite abundant interaction with FcγRIIa and poorly elicited antibody-dependent cellular cytotoxicity, suggesting their Fc features prevent cell killing responses. Restoring FcγRIIIa interaction may be a promising therapeutic approach in HGSOC.

**Highlights:**

- TBAs in ovarian carcinoma patients consistently lack interaction with FcγRIIIa
- Ascites- and serum-derived TBAs have heterogeneous subclass composition
- Systems analysis shows complex serologic differences between TBAs and antiviral responses
- Patient-expressed TBAs demonstrate little antibody-dependent cellular cytotoxicity

## Introduction

Ovarian cancer (OC) accounts for 207,000 deaths annually worldwide, and in the United States, it is estimated to affect 20,890 women and lead to 12,730 deaths in 2025^1,2^. High grade serous ovarian cancer (HGSOC) is the most common and lethal subtype of OC, making up 75% of OC cases and 80% of deaths^1,3^. HGSOC is often diagnosed at an advanced stage and standard first-line treatment consists of surgical cytoreduction and platinum-based chemotherapy. Although most HGSOC patients initially respond well to platinum-based chemotherapy, more than 80% of patients will experience recurrence within three years^4–6^. Recurrent disease is often platinum-resistant and is incurable as there are few alternative treatments. Patients with other cancer types, such as metastatic melanoma and advanced renal-cell carcinoma, have shown responses to and increased survival with immunotherapies^7,8^; however, these therapies, such as checkpoint inhibitors that disrupt immune checkpoint signaling, have shown poor results in HGSOC^9^. Consequently, there is an unmet need for novel immunotherapeutic targets.

In OC, while the adaptive immune response includes both antibody (Ab)- and cell-mediated immunity, substantial barriers in cell-mediated responses limits their therapeutic potential in HGSOC. Tumor-infiltrating B cells (TIL-Bs) have been found in more than 40% of HGSOCs and are strongly correlated with survival^10^; tumor-binding autoantibodies (TBAs) produced by TIL-Bs recognize tumor-associated antigens (TAAs) like mutant p53 and many self-antigens often overexpressed in tumor tissues^10^. TIL-Bs also play a key role in promoting anti-tumor effects by presenting antigens (Ags) to T cells and promoting the T cell response^11^. These T cells include CD4^+^ helper T cells that stimulate immune cell populations, such as macrophages and dendritic cells, and activated CD8^+^ T cells that recognize Ags expressed on the surface of OC cells and subsequently induce tumor cell apoptosis^12–14^. However, while the presence of these T cell populations in the tumor microenvironment (TME) has been associated with improved patient survival in OC^11,13^, progressive T cell exhaustion eventually hinders their anti-tumor responses through persistent Ag-driven increases in inhibitory receptor expression and loss of effector functions. Despite the development of immune checkpoint inhibitors designed to help overcome this exhaustion, HGSOC has been poorly responsive to T cell checkpoint immunotherapies developed to date^15^. TBAs may offer an alternative path to tumor cell killing for cancers such as OC with limited T cell-mediated immunity.

IgG antibodies (Abs) primarily induce tumor cell death by activating cellular processes such as antibody-mediated cellular cytotoxicity (ADCC) and phagocytosis (ADCP). During these responses, Abs first bind to the tumor cell through its fragment antigen-binding (Fab) region, and then bind Fcγ receptors (FcγRs) on immune cells, which interact with the Fc domain of IgG and regulate ADCC, ADCP, and the release of inflammatory factors^16–18^. FcγRs preferentially bind to the IgG subclasses IgG1 and IgG3 over IgG2 and IgG4^19,20^, with these subclass distinctions defined by sequence differences in the Fc region; however, the receptors have an overall low affinity for IgG Abs and rely on the avidity of several Abs bound to a target for activation^21,22^. The interaction between the antibody and FcγRIIa, expressed on macrophages and other myeloid cells, initiates ADCP. Similarly, the interaction between the Ab and FcγRIIIa, found on natural killer (NK) and other innate immune cells, is particularly critical for ADCC and other types of cell killing. For instance, therapeutic monoclonal Abs are routinely engineered to have a higher affinity for FcγRIIIa in order to drive more potent cell killing^23^. In addition to IgG subclass, Ab glycosylation significantly impacts the extent to which an Ab binds to Fc receptors. In fact, IgG Fc can be fucosylated during expression by B and plasma cells, resulting in nearly complete loss of FcγRIIIa binding and, consequently, lower ADCC^24^.

TBAs have been identified as important components of the anti-tumor immune response across multiple cancer types. For example, in lung adenocarcinoma, TBAs contribute to anti-tumor immunity by recognizing endogenous retrovirus proteins, expression of which is reactivated in cancer; this response is enhanced by and contributes to immunotherapy^25^. In murine melanoma models, TBAs are produced as part of a strong humoral response following tumor inoculation and subsequent immunotherapy, and continue to provide protection on subsequent tumor challenges^26^. Despite the widespread existence of TBAs in HGSOC, they are generally not thought to participate strongly in direct cell killing of tumor tissues^27^. Various theories have been proposed for this observation, including the presence of immune checkpoint receptors such as CD47, but these theories have not translated to effective therapies^28–31^. Research has primarily focused on the Fab-mediated Ag-binding properties of TBAs, while the critical Fc-mediated capability of these autoantibodies to simply engage receptors and drive downstream cytotoxic responses has remained unexplored^32^. Given the substantial abundance of TBAs in HGSOC and their cytotoxic potential in other malignancies, understanding why these autoantibodies do not participate strongly in the cytotoxic killing of tumor cells is critical for developing complementary immunotherapies. Activating TBA-mediated effector functions may address current treatment resistance and broaden the spectrum of viable immunotherapies for HGSOC.

We hypothesized that TBAs from HGSOC patients carry features within their Fc domain that prevent effective tumor cell killing. Therefore, using a multiplexed approach to characterize TBA Fc properties, we profiled antibody type, subclass, and interactions with Fc receptors, comparing them to common anti-viral Ab responses in two clinical HGSOC cohorts containing ascites—fluids in close proximity to the TME that accumulate high concentrations of Abs produced by TIL-Bs—and serum samples. We found that, while TBAs were heterogeneously composed of IgG subclasses, there was a consistent and dramatic lack of interaction between TBAs and FcγRIIIa. Consequently, TBAs derived from patients resulted in minimal ADCC responses against tumor cells. These results indicate that, while TBAs may be abundant in most patients, strategies to restore these requisite interactions to drive cell killing will be essential to their therapeutic benefit.

## Results

### Tumor-binding autoantibodies lack interaction with FcγRIIIa

We first sought to confirm the presence of TBAs in HGSOC, consistent with earlier studies^33^. To do so, we stained patient primary solid tumor tissue and ascites samples for IgG and PAX8, a transcription factor highly expressed in OC. Tumor cells derived from these solid tumor tissues and ascites samples were consistently positive for both IgG and PAX8 staining (Figs. 1A–B, Table S1). Quantification of IgG staining revealed that ∼40% of tumor cells from the patient samples exhibited 21–50% positive staining, ∼25% of samples showed >50% positive staining, and the remaining samples showed ≤20% positive staining (Fig. 1C, left). Tumor tissues and ascites were scored between 1^+^–3^+^ by a pathologist based on the intensity of IgG shown: >25% of the samples had a score of 2^+^ and >20% had a score of 3^+^ (Fig. 1C, right).

**Figure 1.**
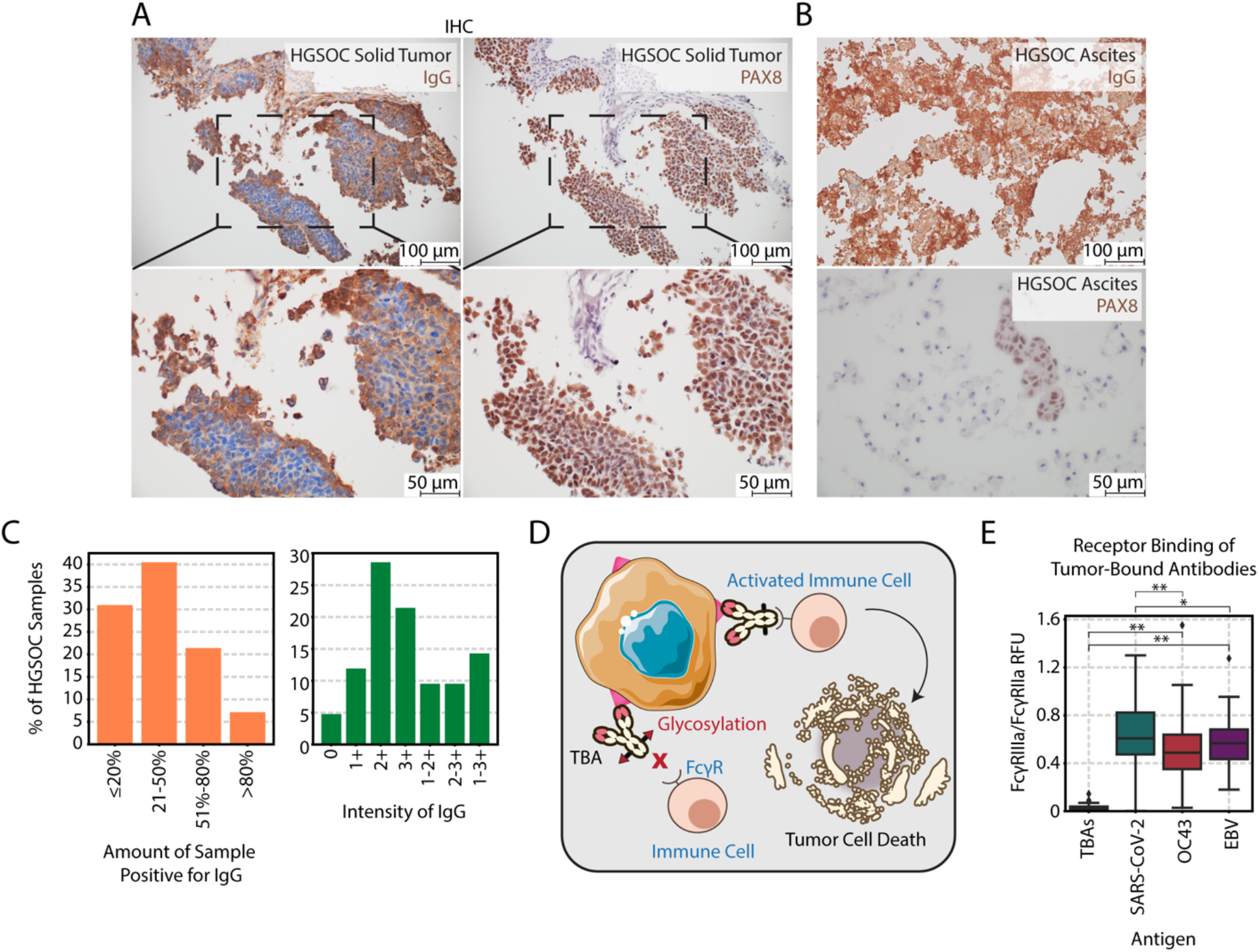
While patients produce endogenous IgG autoantibodies specific to their tumors, systems serology reveals a lack of TBA interaction with FcγRIIIa. A) Anti-human IgG (left) immunohistochemistry (IHC) staining of an HGSOC primary solid tumor specimen from patient 9. Anti-PAX8 staining (right) is included as a tumor-specific lineage marker. Scale bars represent 50 µm or 100 µm, where noted. B) Anti-human IgG (top) IHC staining of an HGSOC ascites specimen from patient 56. Anti-PAX8 staining (bottom) of the same ascites specimen from patient 56. Scale bars represent 50 µm or 100 µm, where noted. C) Quantification of the staining across patients with available samples in the cohort (n=29). Samples with heterogeneous staining were assigned a range instead of a single score. D) Visual depiction of our hypothesis. While TBAs bind to tumor cells, their Fc properties may prevent interaction with the FcγRs on immune cells. E) The ratio of FcγRIIIa to FcγRIIa binding across patients, separated by Ab antigenic target (n=61). Fluorescence is indicated by relative fluorescence units (RFU). In E), significance was determined between the TBAs and anti-viral Abs using a Kruskal-Wallis H Test followed by Dunn’s post-hoc test with Bonferroni corrections. Here, “*” and “**” represent p-values of less than 0.05 and 0.0005, respectively. See also Figure S1 and Table S1.

Given the presence of these autoantibodies and the limited role of TBAs in mediating effector responses, we hypothesized that the requisite Fc interactions needed to direct tumor elimination may be deficient (Fig. 1D). To evaluate this hypothesis, we adapted so-called systems serology assays to characterize TBAs and their Fc interactions in a multiplexed manner, allowing us to simultaneously measure multiple parameters^34–38^. In this systems serology approach, we quantified the relative abundances of TBAs, characterized their IgG subclasses, and linked them to Fc effector functions by measuring their binding to FcγRs. Since the antigen (Ag) targets of these TBAs are likely complex and variable between patients, we isolated the TBAs from patient ascites and serum according to their binding to a well-characterized HGSOC cell line^33,39^, OVCAR3, ensuring consistent and standardized conditions for Ag binding (Fig. S1A). Repeated analysis of the same samples yielded comparable results, demonstrating the assay’s reproducibility with an HGSOC cell line (Figs. S1B–C). Using this assay, we quantified the extent of TBA interaction with FcγRIIa and FcγRIIIa, the key receptors driving ADCP and ADCC. These Fc receptor interactions were then compared to those elicited by several acute infections (SARS-CoV-2, OC43), chronic infections (EBV), or vaccine-induced responses (SARS-CoV-2), as immune responses to these common viruses are well-characterized. TBAs showed an almost complete lack of interaction with FcγRIIIa compared to these viral baselines (Fig. 1E).

### Ascites- and serum-derived autoantibodies have heterogeneous subclass composition and functional interactions

Based on the intriguing differences in Fc receptor interactions between TBAs and the anti-viral serology responses, we sought to more deeply characterize these interactions. The Fc interactions were quantified alongside the abundance of each IgG TBA subclass. TBAs were composed of higher levels of IgG2–IgG4, while in comparison, patient-derived anti-EBV Abs were predominantly IgG1 (Fig. 2A). To evaluate subclass distribution within individual patients, we examined the proportion of total fluorescence attributed to IgG1. We found that TBAs were less IgG1-skewed as compared to anti-EBV Abs, suggesting a more diverse subclass profile in the TBA response (Fig. 2B). Anti-OC43 Abs demonstrated a similar trend, with IgG1 being the predominant subclass, although some patients had elevated levels of anti-OC43 IgG2 (Fig. 2C). Consistent with our anti-EBV Ab findings, TBAs had a reduced IgG1 skew as compared to anti-OC43 Abs, further supporting a broader subclass distribution in the tumor-specific response (Fig. 2D). By contrast, anti-SARS-CoV-2 Abs showed patient-specific variation and were variously skewed toward IgG1 or IgG4 (Fig. 2E). This can be attributed to the timing of our sample collection throughout and following the pandemic; SARS-CoV-2 mRNA vaccine boosting leads to class switching to IgG4^40^ (Supplemental Figs. 2A–D). As expected, examination of total fluorescence confirmed that TBAs demonstrated lower amounts of IgG4 as compared to anti-SARS-CoV-2 Abs derived from our patients (Fig. 2F). TBAs displayed heterogeneous subclass composition, whereas anti-viral Abs skewed more heavily toward IgG1, emphasizing distinct features within each serological response.

**Figure 2.**
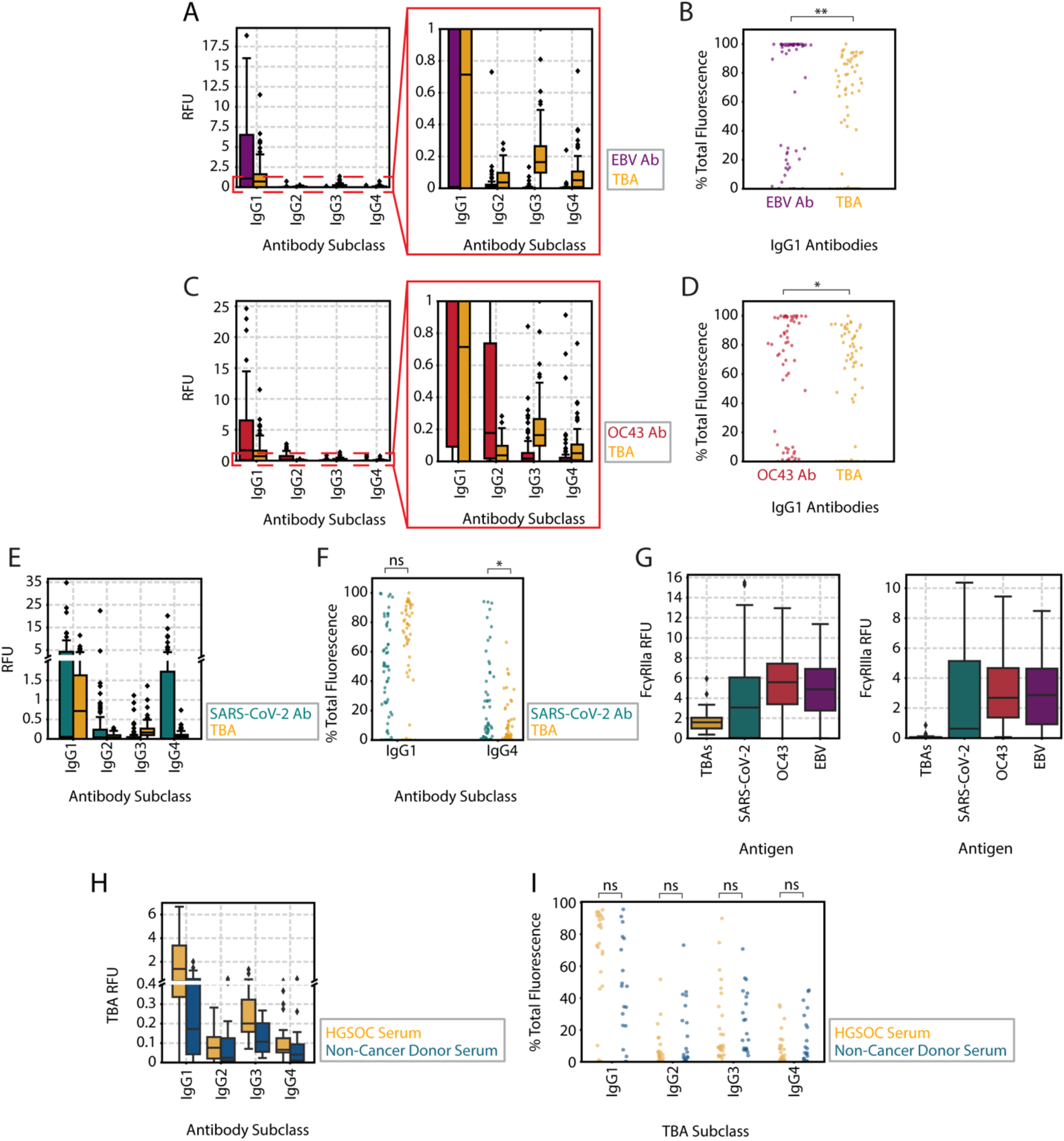
Ascites- and serum-derived Abs have heterogeneous subclass composition and functional interactions. A) IgG subclass compositions for patient anti-EBV Abs as compared to patient TBAs (n=61). B) Percentage of anti-EBV Ab and TBA RFU that is associated with IgG1 normalized to the total IgG signal across all four subclasses (n=61). C) IgG subclass compositions for patient anti-OC43 Abs as compared to TBAs (n=61). D) Percentage of anti-OC43 IgG and TBAs that is associated with IgG1 normalized to the total IgG signal across all four subclasses (n=61). E) IgG subclass compositions for patient anti-SARS-CoV-2 Abs as compared to TBAs (n=61). F) Percentage of anti-SARS-CoV-2 IgG and TBAs that is associated with IgG1 or IgG4, normalized to the total IgG signal across all four subclasses (n=61). G) Quantification of FcγRIIa (left) and FcγRIIIa (right) binding for IgG isolated according to their binding to tumor cells, SARS-CoV-2 Spike, OC43 Spike, or EBV gp350 (n=61). H) IgG subclass quantification for TBAs derived from patient serum (n=26) and non-cancer donor serum samples (n=20). Non-cancerous donor samples were included to provide a baseline for comparison, allowing us to distinguish tumor-associated Ab features from those observed in individuals without cancer. I) Percentage of patient (n=26) and non-cancerous donor serum (n=20) RFU associated with each IgG subclass normalized to the total IgG signal across all four subclasses. In B), D), F), and I), significance was determined using a Mann-Whitney U test to compare TBAs with anti-viral Abs, as well as patient-derived TBAs with those from non-cancerous donor serum. Here, * and ** represent p-values of less than 0.05 and 0.0005, respectively. See also Figure S2.

We next quantified Fc interactions across the Ab groups. We first saw that there was appreciable binding of TBAs to FcγRIIa, indicating a detectable and measurable Ab response (Fig. 2G, left). The amount of TBA binding to FcγRIIa was approximately 27-fold greater than the background fluorescence observed within the assay. However, TBA binding to FcγRIIIa was below the limit of detection, except in a small subset of samples (Fig. 2G, right). Taken together, these findings indicate that the lack of FcγRIIIa engagement is not due to an absence of TBAs, but rather, demonstrates a selective deficit in FcγRIIIa binding.

Given that TBAs may be locally secreted from TIL-B cells or systemically disseminated^27^, we next sought to examine the subclass representation of the TBAs across sample sources. We focused this analysis on serum-derived TBAs, as healthy individuals do not have ascites. In both serum types (patient serum, non-cancerous donor serum), TBAs were skewed toward IgG1 and IgG3 subclasses, with IgG2 and IgG4 being comparatively less abundant (Fig. 2H). Patient serum consistently contained higher amounts of TBAs across all four IgG subclasses in comparison to the OVCAR3-isolated IgG derived from non-cancerous donor serum (Fig. 2H). Notably, IgG1 and IgG3 constituted a greater fraction of total IgG in patient serums, indicating that TBAs in patient serum are biased towards subclasses associated with potent effector functions (Fig. 2I).

To further characterize the TBAs, we then examined their specificity for several common TAAs (p53, matrix metalloproteinase-14 (MMP-14), EGFR, HER2). TBAs targeting p53 were the most abundant across all four IgG subtypes, and these antibodies were skewed toward IgG1 as compared to TBAs targeting MMP-14, EGFR, and HER2 (Fig. S2E). Since MMP-14 was previously identified as a recurrent antigenic target of TBAs^33^, it was unsurprising to see that IgG1 TBAs targeting MMP-14 were the next most abundant group in our cohort (Fig. S2E). Interestingly, of the four tumor-associated Ags profiled amongst the TBAs, those targeting EGFR were uniquely skewed toward IgG4 (Fig. S2E).

### PCA reveals complex serologic differences between TBAs and anti-viral responses

While the initial examination of our dataset established an overview of patterns within the measurements, a formal multivariate analysis was required to uncover how these measurements related to one another in aggregate. We next used principal component analysis (PCA) to quantify multivariate relationships and reveal dominant sources of variation across multiple Ab measurements in our patients (Fig. 3A). This dimensionality reduction approach reveals underlying relationships that are obscured when examining individual Ab measurements separately. By capturing coordinated variation in Fc characteristics and subclass composition, PCA unifies these features into a single framework, providing deeper insight into how they collectively influence effector function and contribute to the impaired effector response observed in the patients. Four principal components (PCs) captured >80% of the variance, representing most of the meaningful variation in the data (Fig. S3A). Patient samples did not separate according to race or treatment history (Figs. S3B–G).

**Figure 3.**
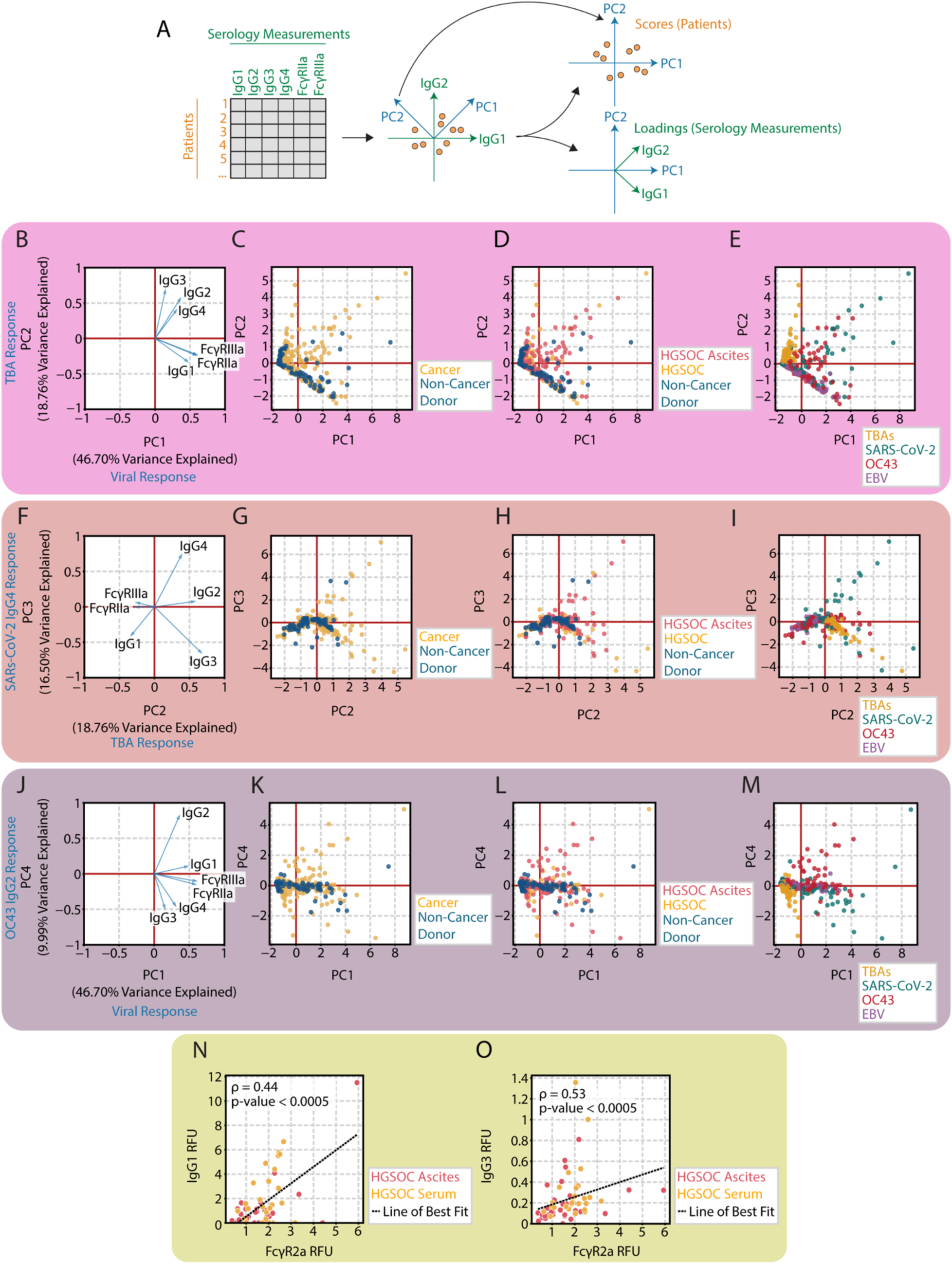
PCA reveals complex serologic differences among TBAs and anti-viral responses. A) Z-scored data were decomposed by PCA into scores (patient samples) and loadings (serology measurements). IgG1 and IgG2 loadings are shown as representative examples of the six serology measurements used in the analysis. B) PCA loadings showing the contribution of each variable to PC1 and PC2. PC1 is positively associated with an increase in IgG1 and FcγR binding. PC2 is positively associated with an increase in IgG3. C–E) Scores plots for PC1 and PC2, colored by patient disease status, sample type, and antigenic target for each measurement (n=81). F) PCA loadings showing the contribution of each variable to PC2 and PC3. PC3 represents strongly IgG4-specific responses, which occurred in many of the anti-SARS-CoV-2 measurements. G–I) Scores plots for PC2 vs. PC3, colored by patient disease status, sample type, and antigenic target for each measurement (n=81). J) PCA loadings showing the contribution of each variable to PC1 and PC4. K–M) Scores plots for PC1 vs. PC4, colored by patient disease status, sample type, and antigenic target for each measurement (n=81). N) Amount of IgG1 and O) IgG3 TBAs derived from patient ascites and serum as a function of FcγRIIa interaction (n=61). In N) and O), Spearman correlation was used to evaluate the relationships between FcγRIIa binding and amounts of IgG1 and IgG3 TBAs, respectively. See also Figure S3.

PCA identified distinct immunological patterns separating TBAs from anti-viral Abs. The loadings, which represent each variable’s contribution to the PCs, showed that IgG1, FcγRIIa, and FcγRIIIa were positively associated with PC1, while IgG2, IgG3, and IgG4 were positively associated with PC2 (Fig. 3B). When examining the scores, which reveal how each observation aligns with the loading patterns, cancer patients were distributed along both PC1 and PC2, but with greater spread along PC2 (Fig. 3C). Ascites samples were more widely distributed across both components than serum samples (Fig. 3D). Notably, TBAs separated distinctly from anti-viral Abs (SARS-CoV-2, OC43, EBV) (Fig. 3E). PC1 was driven primarily by variation in the anti-viral response, while TBAs were characterized by negative PC1 and positive PC2 scores (Fig. 3E). Given the strong positive association between PC1 scores and FcγR binding, the negative PC1 positioning of TBAs indicates minimal FcγR interaction compared to the anti-viral Abs (Fig. 3B, Fig. 3E).

Analysis of PC3 revealed an additional anti-SARS-CoV-2-specific pattern of Ab variation within our patient cohort, with IgG4 subclass as the sole positive contributor to this component (Fig. 3F). PC3 was primarily associated with a subset of cancer patients (Fig. 3G) and SARS-CoV-2 Abs derived from their ascites, which were notably enriched for IgG4 (Fig. 3H–I). The IgG4-skewed response likely reflects SARS-CoV-2 mRNA vaccination boosting; our ascites samples were collected during and post-pandemic, while most of our serum samples were obtained pre-pandemic (Figs. S2A–D). This pattern is consistent with previous observations of IgG1 to IgG4 class switching following two doses of the mRNA vaccine^40^.

PC4 distinguished IgG2 responses from IgG3 and IgG4 responses (Fig. 3J) while also separating cancer samples from non-cancerous donor samples (Fig. 3K). Anti-OC43 Abs derived from ascites had a strong positive association with the IgG2 response associated with PC4 in the positive direction (Fig. 3J, Figs. 3L–M). In contrast, TBAs and anti-SARS-CoV-2 Abs were the predominant contributors to the IgG3 and IgG4 responses (Fig. 3J, Fig. 3M). These findings corroborate our earlier observations of elevated IgG2 anti-OC43 Abs in certain patients, while TBAs and anti-SARS-CoV-2 Abs were more abundant as IgG3 and IgG4, respectively (Fig. 2E, Fig. 2H).

To better understand the mechanisms underlying the distinct Fc receptor binding profiles of the anti-viral Abs versus TBAs, we examined whether subclass composition could account for these differences. As expected, given the well-established roles of IgG1 and IgG3 in potently driving effector responses^41,42^, patients with elevated IgG1 (Fig. 3N) and IgG3 (Fig. 3O) TBAs exhibited increased FcγRIIa binding. However, subclass composition cannot fully explain the TBAs’ loss of FcγRIIIa interaction. If this deficiency were solely attributable to subclass differences, TBAs would be enriched for lower-affinity IgG2 or IgG4 subclasses. Instead, TBAs were predominantly composed of IgG1 and IgG3 (Figs. 2H–I), yet still demonstrated detectable FcγRIIa binding that exceeded their interaction with FcγRIIIa (Fig. 1E, Fig. 2G).

### Patient-expressed TBAs poorly elicit cell-mediated cytotoxicity

Finally, based on our observation that patient TBAs interacted poorly with FcγRIIIa, we hypothesized that they would be ineffective in eliciting ADCC. Therefore, we directly tested their functional capacity. As a basis of comparison, we compared TBA-elicited ADCC to that of Cetuximab, a wild-type IgG1 monoclonal Ab. At the concentrations tested, Cetuximab led to significantly less cell-bound IgG as compared to patient TBAs (Fig. 4A). However, despite markedly lower tumor cell binding than TBAs, cell-bound Cetuximab more readily interacted with FcγRIIIa compared to patient TBAs (Fig. 4B).

**Figure 4.**
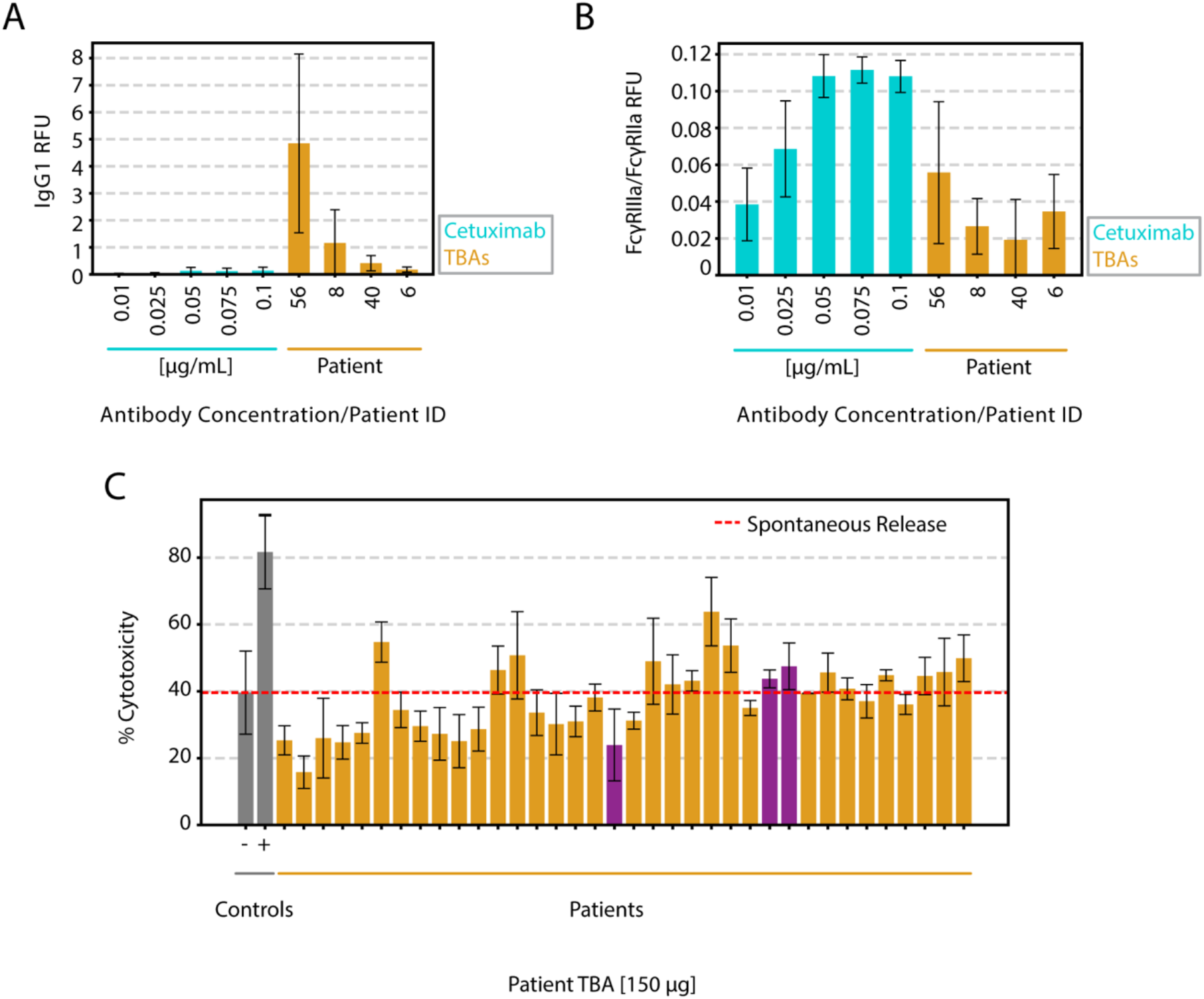
TBAs exhibit diminished capacity for ADCC. A) Quantification of Cetuximab, an IgG1 Ab, binding to OVCAR3 cells as compared to patient TBAs by fluorescence. Cetuximab was incorporated at concentrations of 0.01–0.1 µg/mL. B) Quantification of Cetuximab binding to FcγRIIIa as compared to patient TBAs via fluorescence. C) Percent cytotoxicity of unmodified patient TBAs, quantified by an *in vitro* effector response assay (n=36). All patient TBAs were normalized to the same concentration. Cetuximab was used as a positive control (+). NK cells co-incubated with OVCAR3 cells alone functioned as the negative control (-). A purple bar indicates a patient that was evaluated in A) and B). Data are represented as average ± standard deviation. See also Figure S4.

We then measured ADCC in response to patient TBAs. The majority of the patients exhibited negligible TBA-mediated cytotoxicity, with killing activity at or below the negative control (Fig. 4C). Taken together, while TBAs bound to tumor cells to a greater degree than Cetuximab (Fig. 4A), they interacted comparatively less with FcγRIIIa and poorly elicited ADCC (Figs. 4B–C).

Lastly, we sought to examine the direct impact of glycosylation on Ab-FcR interactions. We suspected fucosylation of the TBAs to be the reason for their poor receptor engagement^43^, so we enzymatically treated isolated TBAs using α1-2,4,6 Fucosidase O (FucO) to remove the core fucose, and an endoglycosidase for specifically cleaving all N-linked glycans (EndoS) (Fig. S4A). However, the core fucose in Abs is attached to the innermost GlcNAc residue, creating significant steric hindrance that shields the fucose from fucosidase hydrolysis^44^. As expected, FcγRIIa and FcγRIIIa binding quantification pre- and post-treatment revealed that while TBA binding with FcγRIIa increased slightly with only FucO, there was no significant difference in FcγRIIIa binding (Fig. S4B). Treatment with either EndoS or a combination of FucO with EndoS drastically reduced interaction of TBAs to both FcγRs, demonstrating that Ab-immune cell receptor interaction was glycosylation dependent. Two other fucosidases, FucA1 and BfFucH, had previously demonstrated some activity in removing core fucose from intact Abs, with varying success^44,45^. However, treatment of isolated TBAs with either FucA1 or BfFucH did not yield significant changes in receptor binding (Figs. S4C–E). Ultimately, our finding that these enzymatic strategies are ineffective against the complex Fc glycan is consistent with prior literature^45^.

## Discussion

Here, we confirmed, as has been observed elsewhere^46–50^, that IgG TBAs are widespread and abundant in OC patients. Although previous work in this field has focused on the origin and the antigenic targets of TBAs^33^, IgG Fc domain interactions are crucial for inducing cytotoxic killing of cells through interaction with FcγRs^51–54^. Hypothesizing that these Abs may poorly elicit cell killing because of their Fc properties, we adapted the toolkit of systems serology assays to characterize these TBAs^34–38,55^. We used these tools to comprehensively profile patient TBA Ag -specificity, subclass composition, and binding to Fc receptors. The development of our novel cell-based Ab pulldown assay for TBA isolation takes advantage of an OC cell line to selectively draw out TBAs, circumventing the challenges of Ag heterogeneity within patients and frequent absence of known antigenic targets^25,27,33,55,56^. Anti-viral Abs from the patients were predominantly made up of IgG1, while TBAs were heterogeneously comprised of both IgG1 and IgG3. While the subclass composition and abundance of TBAs was heterogeneous, they consistently and dramatically lacked interaction with FcγRIIIa compared to anti-viral IgG. Correspondingly, TBAs elicited little to no cell killing. Ab responses are a critical defense mechanism in cancer, but our work reveals that despite selectively recognizing the tumor, TBAs are unable to elicit successful downstream cytotoxic killing of tumor cells because they lack necessary interactions within the Fc domain, allowing tumors to evade cell-mediated Ab immunity.

The specific underlying regulatory mechanisms that govern natural fucosylation and broader glycosylation patterns in Abs remains incompletely understood. In general, fucosylation is known to be modulated in an Ag-specific manner, alongside contextual cues. Evidence in acute infection has shown that cell membrane- and viral envelope-associated Ags drive the production of afucosylated IgG, while IgG against internal Ags demonstrate higher fucosylation levels^38,57–63^. The vast majority of overall IgG in healthy individuals is fucosylated, which is thought to serve as a protective measure against harmful attack on healthy tissue^64,65^. Increased levels of core fucosylation, particularly in OC, have been observed in tumor tissues compared to normal tissues, suggesting specific regulatory mechanisms participating in tumor progression^66,67^. Some autoimmune diseases, such as rheumatoid arthritis, are also associated with elevated levels of IgG fucosylation, suggesting that there might be general regulation of fucosylation as feedback after chronic autoimmunity^68,69^. However, our data suggest divergent fucosylation exists between TBAs found on the cell surface and IgG against viral Ags within the same patient, consistent with Ag-specific regulation, as evidenced by their distinct FcγR binding profiles. Our findings provide compelling evidence that the diminished binding of FcγRIIIa binding observed in TBAs, compared to Cetuximab, is driven by differences in Fc domain glycosylation. While tumor cell binding in TBAs was not impaired, indicating intact Fab-mediated Ag recognition, fucosylation selectively disrupted TBA FcγRIIIa engagement. These findings support our focus on Fc-specific mechanisms as the primary contributor the impaired effector function of TBAs. A better understanding of the regulatory events driving TBA fucosylation would in turn suggest targetable mechanisms of restoring tumor cell killing.

Alternative strategies for the selective removal of core fucose residues from TBAs involve significant experimental complexities that present difficulties in translation to clinical applications. Current afucosylation approaches, including trifluoroacetic acid treatment, engineered cell lines with reduced fructosyltransferase expression, and targeted gene silencing, involve complex, time-intensive protocols that are unideal for a clinical setting^70–75^. Our difficulty in restoring TBA cytotoxic capabilities following enzymatic modification further demonstrates the complexity of the issue. Fucosylation is a critical barrier to effective receptor engagement of the TBAs. Taken together, these underscore the urgent need for developing clinically viable strategies to address TBA fucosylation in OC.

The objective of this work was to study the body’s natural immune response against OC. Our work aimed to establish a foundation for determining whether TBA modification might represent a viable therapeutic strategy for HGSOC and potentially other cancers, in addition to providing insights into the regulation of Ab-mediated immunity in the TME. Understanding the mechanisms by which TBAs engage with, or fail to engage with, other components of the immune system is critical for informing the development of more effective strategies for harnessing the humoral immune response against ovarian cancer and potentially other malignancies^27^. Novel therapeutic opportunities that exploit Ab effector functions to enhance anti-tumor responses could be revealed through this work^33,76,77^. Ultimately, our research highlights the potential for a fundamentally new approach to cancer immunotherapy by harnessing and enhancing naturally occurring anti-tumor Ab responses, leading to improved survival rates and enhanced quality of life for patients.

### Limitations of the study

While we were able to systematically profile TBAs in HGSOC, our work is limited by modest sample size and scope limited to a single cancer type. Given that HGSOC treatment and outcomes are relatively homogenous compared to other cancer types, our focus precludes analysis of how TBA Fc features might change with immunotherapy treatment or associate with distinct long-term outcomes^25,78,79^. Expanding the scope and size of our clinical cohort, including across cancer types, would provide additional data from which we would identify clinical associations that may have been missed in our smaller cohort. It might also reveal mechanistic determinants of Fc features, such as if these change with therapies, like checkpoint inhibitor treatment or with certain cancer types. Finally, translating our findings into clinical applications remains challenging due to limitations in HGSOC humanized mouse models. However, humanized patient-derived xenograft models would offer a promising approach for investigating effective TBA reactivation strategies^80–82^.

## Resource availability

### Lead contact

- Requests for further information and resources should be directed to and will be fulfilled by the lead contact, Aaron S. Meyer.

### Materials availability

- This study did not generate new unique reagents.

### Data and code availability

- The raw Luminex and ADCC data are available upon request from the lead contact. All clinical parameters for all two cohorts are reported in Table S2. Due to study participant confidentiality concerns, per patient clinical data cannot be publicly released.
- This paper does not report an original code.
- Any additional information required to reanalyze the data reported in this work is available from the lead contact upon request.

## Acknowledgements

This study was supported by the Emerging Leader Award from the Mark Foundation for Cancer Research, the Predoctoral Fellowship Award from the Jonsson Comprehensive Cancer Center, and the National Institutes for Health research grant R01CA240965 (PKK). The authors gratefully acknowledge the patients who consented for their ascites samples to be collected by the University of Wisconsin Carbone Cancer Center BioBank, supported by P30 CA014520, and Drs. Hannah Micek and Ning Yang for processing the samples. S.M., G.A.D., and L.B.J., are partially supported by research funding provided by the Department of Veteran Affairs (grant no. I01BX006019 and I01BX006411 to S.M.). The authors also gratefully acknowledge the UCLA/CFAR Virology Core Lab, supported by 5P30 AI028697.

## Author contributions

M.L. designed and conducted the experiments, performed data analysis, and wrote the manuscript. M.T. performed the *in vitro* effector response assays. L.B.J. designed and conducted the immunohistochemistry staining of patient samples, contributed to the manuscript, and advised throughout the study. S.D.T. designed and contributed to pilot experiments for the cell-based Ab pulldown and Ag-specific Ab analysis of patient TBAs. H.D. and A.B. contributed to experiments, including the Ab-specific antibody analysis. G.A.D. and L.B.J. processed the patient samples. S.M., L.B.J., and G.A.D. determined clinical characteristics and demographic information of patient samples. Y.T. and J.G. contributed to the immunohistochemistry staining of patient samples. N.M provided pathological scoring of the immunohistochemistry patient samples. P.K.K. procured patient samples. S.M. provided patient samples, provided critical feedback, and advised throughout the study. A.S.M. conceived and supervised the study, designed experiments, and wrote the manuscript. All authors discussed the results and contributed to the final manuscript.

## STAR★METHODS

### Key resources table

| REAGENT OR RESOURCE | SOURCE | IDENTIFIER |
| --- | --- | --- |
| **Antibodies** |  |  |
| Goat F(ab')<sub>2</sub> Anti-Human IgG, Mouse ads-UNLB | SouthernBiotech | Cat# 2043-01;<br>RRID: AB_2795665 |
| Mouse Anti-Human IgG1 Hinge-BIOT (4E3) | SouthernBiotech | Cat# 9052-08;<br>RRID: AB_2796620 |
| Mouse Anti-Human IgG2 Fc-PE (HP6002) | SouthernBiotech | Cat# 9070-09;<br>RRID: AB_2796639 |
| Mouse Anti-Human IgG3 Hinge-PE (HP6050) | SouthernBiotech | Cat# 9210-09;<br>RRID: AB_2796701 |
| Mouse Anti-Human IgG4 pFc'-PE (HP6023) | SouthernBiotech | Cat# 9190-09;<br>RRID: AB_2796686 |
| PAX-8 Mouse Monoclonal Antibody | Cell Marque | Cat# 363M-15 |
| IgG Antibody | Dako | Cat# GA512 |
| **Bacterial and virus strains** |  |  |
| BL21 (DE3) Competent Cells | Thermo Scientific | Cat #EC0114 |
| **Biological samples** |  |  |
| Human peripheral blood mononuclear cells | UCLA/CFAR Virology<br>Core Laboratory | N/A |
| Human serum samples from female donors | BioChemed | Cat#<br>MT050923UC1 |
| Human HGSOC ascites | UCLA Health and<br>University of<br>Wisconsin Carbone<br>Cancer Center<br>BioBank | N/A |
| Human HGSOC serum | UCLA Health | N/A |
| Human HGSOC primary tumor tissue | UCLA Health | N/A |
| **Chemicals, peptides, and recombinant proteins** |  |  |
| SARS-CoV-2 B.1.1.529 (Omicron) S1+S2 trimer<br>protein | Sino Biological | Cat# 40589-V08H26 |
| Human coronavirus (HCoV-OC43) spike protein (S1+S2 ECD) | Sino Biological | Cat# 40607-V08B |
| Recombinant Human MMP-14/MT1-MMP protein | R&D Systems | Cat# 918-MPN |
| Recombinant Human EGFR Protein (ECD, His Tag) | Sino Biological | Cat# 10001-H08H |
| Recombinant Human Her2/ERBB2 Protein (ECD, His Tag) | Sino Biological | Cat# 10004-H08H |
| Epstein-Barr virus (Herpesvirus 4), His Tag | ACROBiosystems | Cat# GP0-E52H6 |
| Recombinant Human His6- (DYKDDDDK)-p53 Protein, CF | R&D Systems | Cat# SP-452 |
| Recombinant Human Fc gamma RIIA/CD32a Avi His Protein, CF | R&D Systems | Cat# AVI1330 |
| Recombinant Human Fc gamma RIIIA (F176) His Avi Protein, CF | R&D Systems | Cat# AVI8894 |
| Streptavidin-PE | Bio-Rad | Cat# 171304501 |
| Sodium pyruvate | Cytiva | Cat# SH30239.01 |
| Phosphate buffered saline | Fisher Scientific | Cat# BP3991 |
| Bovine serum albumin | Sigma-Aldrich | Cat# H6512 |
| Penicillin-streptomycin solution | Cytiva | Cat# SV30010 |
| FucA1 | R&D Systems | Cat# 7039-GH-010 |
| α1-2,4,6 Fucosidase O | New England Biolabs | Cat# P0749S |
| SOC Outgrowth Medium | New England Biolabs | Cat# B9020S |
| Kanamycin sulfate | Sigma-Aldrich | Cat# K1377-5G |
| LB Broth | Sigma-Aldrich | Cat# L3022-250G |
| Triton X | Sigma-Aldrich | Cat# X100 |
| Isoproyl D thigalactopyranoside | A.G. Scientific | Cat# I-1312-1GM |
| Glycobuffer | New England Biolabs | Cat# B1727S |
| Endo S | New England Biolabs | Cat# P0741S |
| **Critical commercial assays** |  |  |
| DELFI A EuTDA Cytotoxicity Detection Kit | Revvity | Cat# AD0116 |
| EasySep Human NK Cell Isolation Kit | STEMCELL Technologies | Cat# 17955 |
| Melon Gel IgG Purification Kit | Thermo Scientific | Cat# 45212 |
| HistoGel Specimen Processing Gel | Fisher Scientific | Cat# 22-110-678 |
| Pierce BCA Protein Assay Kit | Thermo Scientific | Cat# 23227 |
| Protease Inhibitor Cocktail | Thermo Scientific | Cat# PI78430 |
| Phosphatase Inhibitor Cocktail | Boston Bioproducts | Cat# BP-480-1ml |
| **Experimental models: Cell lines** |  |  |
| Human: OVCAR-3 cells | ATCC | HTB-161 |
| **Recombinant DNA** |  |  |
| pET-28a(+)-BfFucH | This paper | N/A |
| **Software and algorithms** |  |  |
| Scikit-learn | Pedregosa et al.<sup>83</sup> | https://scikit-learn.org/stable/index.html |
| **Other** |  |  |
| Bio-Plex Pro Magnetic COOH Beads | Bio-Rad | Cat# MC100 |
| BD Vacutainer Serum Tubes | BD | Cat# 367815 |
| ProxiPlate 96-shallow well, white opaque | Revvity | Cat# 6006290 |

## Study participant details

Information about study participants can be found in Table S3.

## Method details

### Patient sample collection and processing

Primary HGSOC samples provided by S.M. were collected from patients who provided informed consent through IRB protocols (IRB#10-0727, IRB#22-1970) approved by the UCLA Office of the Human Research Protection Program. Clinical information was obtained from the medical records. p53 status was determined by review of clinical data. Ascites fluid samples were collected from the clinic and immediately processed in the lab. Ascites samples were centrifuged to separate cells from the fluid portion. This fluid portion was then frozen in liquid nitrogen until required for experimental use. Serum was isolated from patient blood samples collected in a vacutainer serum tube (BD, #367815), centrifuged, flash frozen, and stored in liquid nitrogen.

Ascites samples provided by P.K.K. were collected during debulking surgeries of patients diagnosed with Stage III/IV HGSOC at the University of Wisconsin Hospitals and Clinics. The University of Wisconsin Carbone Cancer Center Translational Science BioCore acted as an honest broker under IRB #2016-0934, obtaining informed consent from all participants and de-identifying samples before providing to the research team. The fluid portion of the ascites was isolated by centrifuging at 300 g for 5 minutes and then frozen and stored at -80℃.

See also Table S3.

### Cell-based antibody pulldown and TBA isolation

OVCAR3 cells (ATCC HTB-161) were grown in complete growth media. Patient ascites and serum were prepared by centrifugation at 12,000 rpm for 10 minutes at 4℃, followed by 1:1 dilution with assay buffer (1% bovine serum albumin (BSA) in phosphate-buffered saline (PBS)). The cells were harvested and resuspended using the diluted patient ascites or serum at a concentration of 12.5 µL/125,000 cells, then added to a 96-well round-bottom non-tissue culture treated plate to be incubated for 60 minutes at 4℃, gently shaking.

After the incubation, 100 µL of PBS was added to each well of the plate, followed by centrifugation of the plate at 500 g for 5 minutes at 4℃. The supernatant in each well was removed. The cell pellets were washed by resuspending in PBS. The centrifugation and PBS wash steps were repeated for a total of 7 times. Following the final wash, each cell pellet was resuspended in 50 µL of an NP40 lysis buffer mixture with phosphatase and protease inhibitors and incubated on ice for 5–10 minutes, then spun down at 12,000 rpm for 10 minutes at 4℃. The supernatants were collected and stored as lysates. These lysates containing the isolated TBAs were further diluted with 50 µL of assay buffer prior to long-term storage at -80℃ or immediate use.

### Antigen-specific antibody analysis

A 384-well plate was washed with 1x tris-buffered saline containing 0.1% Tween (TBS-T) prior to use. The capture bead sets for the assay were generated by conjugating anti-pan-IgG, SARS-CoV-2 B.1.1.529 (Omicron) S1+S2 trimer, human coronavirus (HCoV-OC43) spike, MM-P14, EGFR, HER2, Epstein-Barr virus (Herpesvirus 4) gp350, and p53 Abs/proteins to magnetic beads (Bio-Rad).

Beads of each type were combined into a single solution in assay buffer. The bead-assay buffer solution was then added to each well of the plate and washed for a second time. Lysate containing the isolated TBAs from ascites and serum was then added to each well of the washed plate, and co-incubated with the capture beads overnight at 4℃ in the dark, gently shaking. Following the overnight incubation period, the plate was washed again. Alternatively, ascites and serum were directly co-incubated with the capture beads following a 100-fold serial dilution to enable collection of data pertaining to the viral Ags.

Detection-assay buffer solutions were then added to each corresponding well. Biotinylated IgG1 was diluted 1:1000, while phycoerythrin (PE)-conjugated IgG2, IgG3, IgG4 were diluted 1:200. FcγRIIa and FcγRIIIa proteins were diluted 1:200 and separately incubated with streptavidin-phycoerythrin (SAPE) at a 4:1 molar ratio for 15 minutes at room temperature (RT) in the dark prior to addition to the sample plate for tetramerization. The plate was then incubated for 1 hour at RT in the dark, gently shaking. Afterwards, the plate was washed again. For sample wells receiving biotinylated IgG1 detection, an additional incubation period with SAPE-assay buffer solution for 15 minutes at RT was included. The SAPE-assay buffer solution was generated by diluting SAPE 100-fold in assay buffer. The plate was washed a final time, then assay buffer was added to the plate prior to analysis on a Luminex xMAP INTELLIFLEX System.

### Immunohistochemistry staining

Tumor cells isolated from ascites fluid were suspended in HistoGel Specimen Processing Gel prior to preparation in formalin fixed paraffin embedded (FFPE) blocks. Harvested solid tumors were fixed in 10% formalin. Tumor cells and solid tumor specimens were prepared in FFPE blocks. 4 µm sections were deparaffinized, rehydrated, and Ag retrieval was performed in Tris-EDTA. Sections were incubated in blocking buffer and stained with the following antibodies: PAX8 (Cell Marque #363M-15) and IgG (Dako #GA512). Staining was visualized with 3,3-diaminobenzidine (DAB) and counterstained with hematoxylin. For PAX8, fallopian tube was used as a control; secretory epithelial cells stained positive and ciliated epithelial cells stained negative. For IgG, human tonsil was used as a positive control and OVCAR3 cells were used as a negative control.

### In vitro effector response assays

IgG autoantibodies from ascites were purified using the Melon Gel IgG Purification Kit (ThermoFisher #45212) according to manufacturer instructions, to prevent the influence of other factors, such as cytokines, in the functional response measurements. Fresh, unfrozen peripheral blood mononuclear cells (PBMCs) from healthy donor blood were isolated by density gradient centrifugation. Natural killer (NK) cells were isolated from the PBMCs using the EasySep Human NK Cell Isolation Kits (STEMCELL Technologies #17955).

The ADCC activity of TBAs was quantified using the DELFIA EuTDA Cytotoxicity Detection Kit (Revvity #AD0116). Isolated NK cells, OVCAR3 cells, and treated TBAs were co-incubated at a 10:1 effector-to-target ratio, and the DELFIA assay was carried out according to manufacturer instructions using a 96-well microplate (Revvity #6005290) and 2% Triton X as lysis buffer. A Perkin Elmer Victor 3V plate reader was used to obtain time-resolved fluorescence measurements. Analysis was performed using the following formula, where *maximum lysis* is the measurement obtained from incubating BATDA-labeled OVCAR3 cells with lysis buffer, and *spontaneous lysis* is the measurement obtained from only BATDA-labeled OVCAR3 cells:

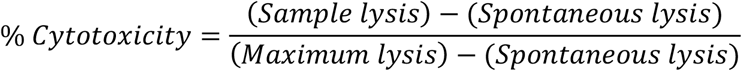

### Expression and isolation of α-L-Fucosidase (BfFucH)

Recombinant BfFucH was cloned into pET-28(+) and transformed into BL21 *Escherichia coli* (BL21 *E. coli*). Briefly, the vector was incubated with *E. coli BL21* on ice for 30 minutes, then heat-shocked for 30 seconds at 42℃. 250 µL of Super Optimal broth with Catabolite repression (S.O.C.) medium was added to the mixture, then shaken at 225 rpm for 1 hour at 37℃. Cells from the transformation reaction were spread on lysogeny broth (LB) plates and incubated overnight at 37℃. The next day, single colonies of transformants were selected and inoculated into LB-medium-kanamycin (50 µg/mL) and cultures were grown overnight at 37℃.

The overnight cultures were inoculated in prewarmed LB medium containing kanamycin once more and then incubated for 1 hour at 37℃, shaking at 250 rpm until confluency. To induce expression, isopropyl β-D-1-thiogalactopyranoside (IPTG) was added to the induced cultures with a final concentration of 0.2 mM. Cultures were grown overnight at 16℃.

Cells were harvested by centrifugation at 15,000 g for 1 minute, then lysed using the freeze-thaw method. BfFucH was then purified using Ni-NTA immobilized affinity chromatography. The purified BfFucH was then analyzed via SDS-PAGE.

### Enzymatic modification of patient TBAs

#### BfFucH treatment

Adapted from Tsai et al.^44^, 10 µg of Melon Gel-isolated patient TBAs were co-incubated at 37℃ overnight with sodium phosphate buffer (pH of 7.5), DI H_2_O, and 10 units of BfFucH for a final reaction volume of 31 µL. Following treatment, FcγRIIa and FcγRIIIa binding were measured by Luminex as described previously using pan-IgG magnetic capture beads.

#### FucA1 (R&D Systems 7039-GH-010) treatment

Adapted from Prabhu et al.^45^, Melon Gel-isolated patient TBAs (0.1 µg/µL) were co-incubated at 37℃ for 7 days with FucA1 (0.1 µg/µL) for a final reaction volume of 20 µL. On day 7, FcγRIIa and FcγRIIIa binding were measured by Luminex.

#### α1-2,4,6 Fucosidase O (FucO) (New England Biolabs #P0749S)

10 µg of Melon Gel-isolated patient TBAs were co-incubated at 37℃ overnight with FucO, Glycobuffer (New England Biolabs #B1727S), with or without Endo S (New England Biolabs #P0741S), and DI H_2_O as necessary for a final reaction volume of 11–12 µL according to manufacturer instructions. Following treatment, FcγRIIa and FcγRIIIa binding were measured by Luminex.

### Principal component analysis

PCA was performed on measurements of IgG1, IgG2, IgG3, IgG4, FcγRIIa, and FcγRIIIa for both patient cohorts. All features were standardized to zero mean and unit variance using the StandardScaler function from *scikit-learn* prior to the analysis. PCA was subsequently applied to the standardized dataset using *scikit-learn*’s PCA implementation.

