## Supplementary Information for "Systems serology of responses against tumor antigens in ovarian cancer reveal disrupted Fc-mediated immunity"

### Supplementary Materials

**Table S1. IgG scoring for immunohistochemistry staining of patient tumor cells and tissues, related to Figure 1.**
Intensity ranged from 1-3 and % positive was based on amount of staining present in each individual sample. A range of "1-2" indicates that regions of the tumor scored as intensities of both 1 and 2.

[a]Longitudinal collections from the same patient; a chemonaive ascites sample was obtained initially (55), and a chemo-treated ascites sample was obtained later.

[b]Longitudinal collections from the same patient; a chemonaive ascites sample was obtained initially (56), and chemo-treated ascites and serum samples were obtained at a later time point.

| Patient Reference Number | Solid Tumor | | Ascites | |
|---|---|---|---|---|
| | % Positive | Intensity | % Positive | Intensity |
| 1 | 40% | 1+ | 70% | 1+ |
| 2 | N/A | N/A | 90% | 1+ |
| 3 | 40% | 1-3+ | N/A | N/A |
| 4 | 40% | 3+ | N/A | N/A |
| 5 | 40% | 2-3+ | N/A | N/A |
| 6 | 40% | 3+ | N/A | N/A |
| 7 | 40% | 2+ | N/A | N/A |
| 8 | 40% | 2-3+ | N/A | N/A |
| 9 | 10% | 3+ | N/A | N/A |
| 10 | 20% | 2+ | N/A | N/A |
| 11 | 10% | 1-3+ | N/A | N/A |
| 12 | 60% | 2+ | N/A | N/A |
| 13 | 30% | 2+ | N/A | N/A |
| 14 | 10% | 1-3+ | N/A | N/A |
| 15 | 30% | 2+ | N/A | N/A |
| 16 | 10% | 1-2+ | N/A | N/A |
| 17 | 40% | 1-3+ | N/A | N/A |

| | | | | |
|---|---|---|---|---|
| 18 | 10% | 2+ | N/A | N/A |
| 19 | 30% | 1+ | N/A | N/A |
| 20 | 5% | 2+ | N/A | N/A |
| 21 | 30% | 1-2+ | N/A | N/A |
| 22 | 60% | 1+ | N/A | N/A |
| 23 | 30% | 2+ | N/A | N/A |
| 24 | 10% | 2+ | N/A | N/A |
| 25 | 20% | 1-2+ | N/A | N/A |
| 55 | 85% | 1-3+ | 80% | 3+ |
| 55[a] | N/A | N/A | 70% | 3+ |
| 56 | N/A | N/A | 80% | 3+ |
| 56[b] | 60% | 1-3+ | 100% | 1-2+ |

**Table S2. Construct and sequence information for BfFucH**. Sequence of the cloned insert gene prior to assembly into the expression construct.

| | |
|---|---|
| Insert length | 1248 |
| Insert sequence | CAACAGAAATACCAGCCTACCGAAGCGAATCTTAAGGCGCGTAGCGAGT<br>TCCAAGATAACAAATTCGGCATCTTTCTCCACTGGGGTCTTTACGCGATGTT<br>AGCAACAGGTGAATGGACCATGACTAATAACAATTTAAACTACAAGGAATA<br>CGCGAAGTTGGCAGGTGGTTTCTACCCATCAAAATTCGACGCGGATAAGT<br>GGGTTGCGGCAATTAAAGCAAGCGGGGCCAAGTACATATGTTTTACGACC<br>CGGCACCATGAAGGCTTTTCTATGTTTGACACAAAATATAGTGACTACAATA<br>TCGTTAAGGCCACACCATTTAAGCGCGACGTCGTCAAAGAATTAGCAGAC<br>GCGTGCGCTAAGCACGGAATTAAGCTGCACTTTTACTACTCCCACATCGA<br>TTGGTACCGCGAGGACGCGCCCCAAGGCCGGACAGGGAGACGGACAG<br>GCCGCCCAAACCCTAAGGGTGACTGGAAATCATACTACCAATTCATGAAC<br>AACCAATTGACCGAACTTTTAACAAACTACGGACCTATAGGTGCCATTTGG<br>TTCGATGGGTGGTGGGACCAAGATATTAACCCTGACTTTGACTGGGAGCT<br>CCCTGAACAGTACGCCCTCATCCACAGATTACAACCCGCATGTCTCGTC<br>GGAAATAATCATCACCAAACCCCCTTCGCAGGTGAGGACATCCAGATATT<br>TGAACGGGACTTACCTGGGGAAAACACAGCCGGCCTGTCAGGTCAGTCA |

| | |
|---|---|
| | GTTAGTCATCTCCCCCTGGAGACCTGCGAAACAATGAACGGCATGTGGG<br>GTTACAAAATCACTGACCAAAACTACAAGTCAACAAAGACATTAATCCATTA<br>CTTGGTCAAGGCAGCGGGCAAGGACGCCAACCTTCTGATGAATATTGGA<br>CCTCAGCCGGACGGAGAGCTCCCTGAGGTTGCTGTACAACGACTTAAAG<br>AGGTTGGTGAATGGATGAGCAAGTACGGCGAGACAATATACGGGACGCG<br>CGGTGGCCTCGTGGCGCCCCACGACTGGGGCGTTACTACCCAGAAGGG<br>AAACAAGCTGTACGTTCATATCCTCAACTTGCAAGACAAAGCGCTTTTCTTA<br>CCGATCGTGGACAAGAAGGTTAAGAAAGCCGTCGTATTCGCCGATAAGAC<br>CCCTGTTCGCTTTACCAAGAACAAAGAGGGGATCGTGTTAGAGCTCGCAA<br>AGGTGCCTACCGACGTGGATTATGTTGTTGAGTTGACGATAGATTAGTAG |

**Table S3. Patient characteristics, related to study participant details, provided as an Excel file.**

Ascites samples were obtained from consented patients at UCLA Health (n=17/35) and from the University of Wisconsin Carbone Cancer Center BioBank (n=18/35). Serum samples were obtained from consented patients at UCLA Health center (n=26).

[a]Longitudinal collections from the same patient; a chemonaive ascites sample was obtained initially and a chemo-treated ascites sample was obtained at a later time point.

[b]Longitudinal collections from the same patient; a chemonaive ascites sample was obtained initially and chemo-treated ascites and serum samples were obtained at a later time point.

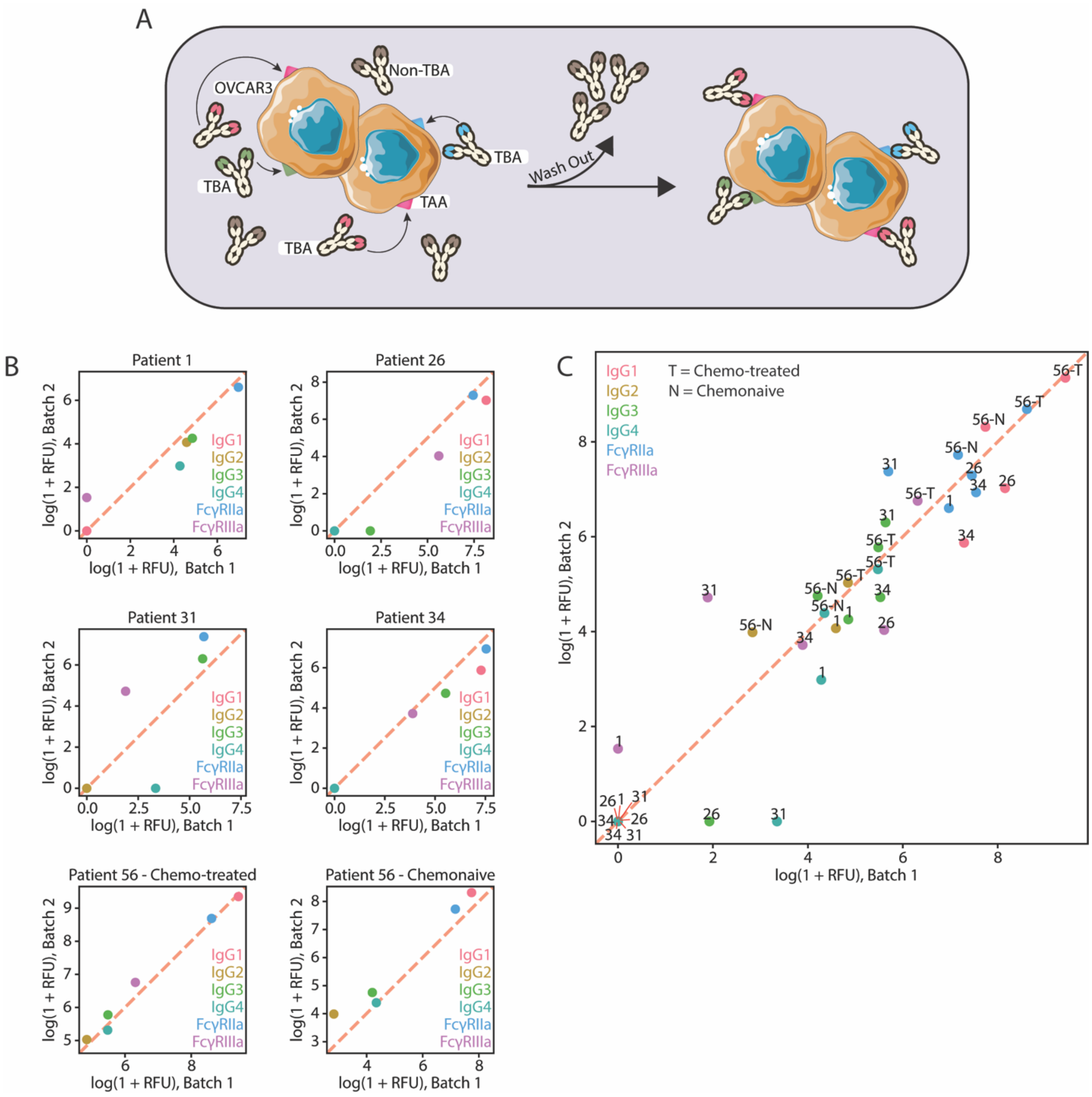


**Figure S1. Reproducibility analysis of TBA subclass composition and functional interactions data demonstrates strong consistency across two experimental batches, related to Figure 1.**

A) Cell-based Ab pulldown and Ab isolation. TBAs from patient samples were isolated based on their binding to OVCAR3 cells.

B) The log(1+RFU)-transformed measurements of TBA subclass and Fc receptor interactions grouped by patient sample and C) displayed across all patients (n=6). FcγRIIIa measurements were not available for patient 56 – chemonaive.

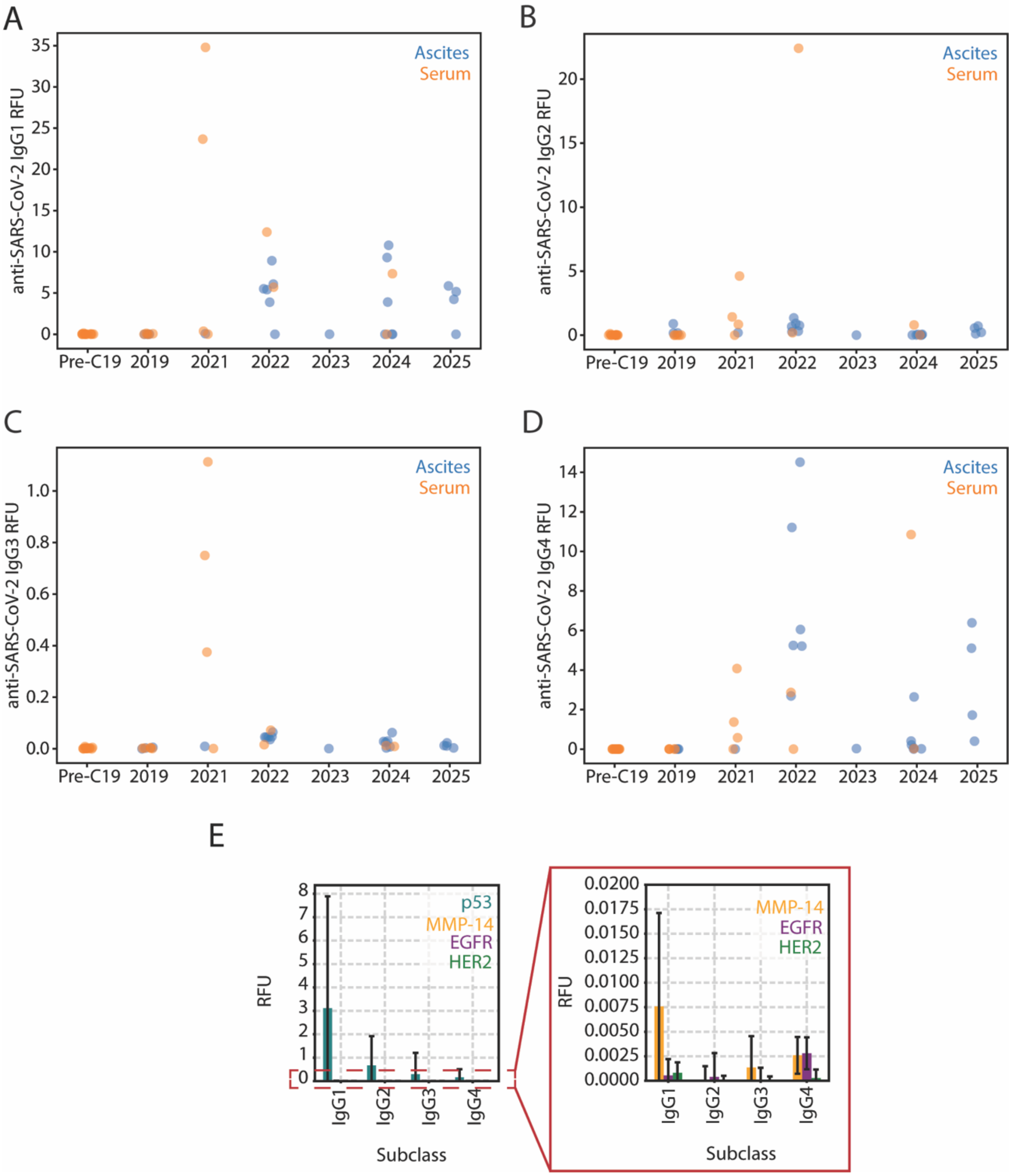


**Figure S2. SARS-CoV-2 vaccination effect and TBA Ag-specificity, related to Figure 2.**

A) IgG1, B) IgG2, C) IgG3, and D) IgG4 anti-SARS-CoV-2 Ab response from patients over duration of sample collection period (n=61).

E) TBA Ag-specificity from patient ascites and serum (n=48). Data are represented as average ± standard deviation.

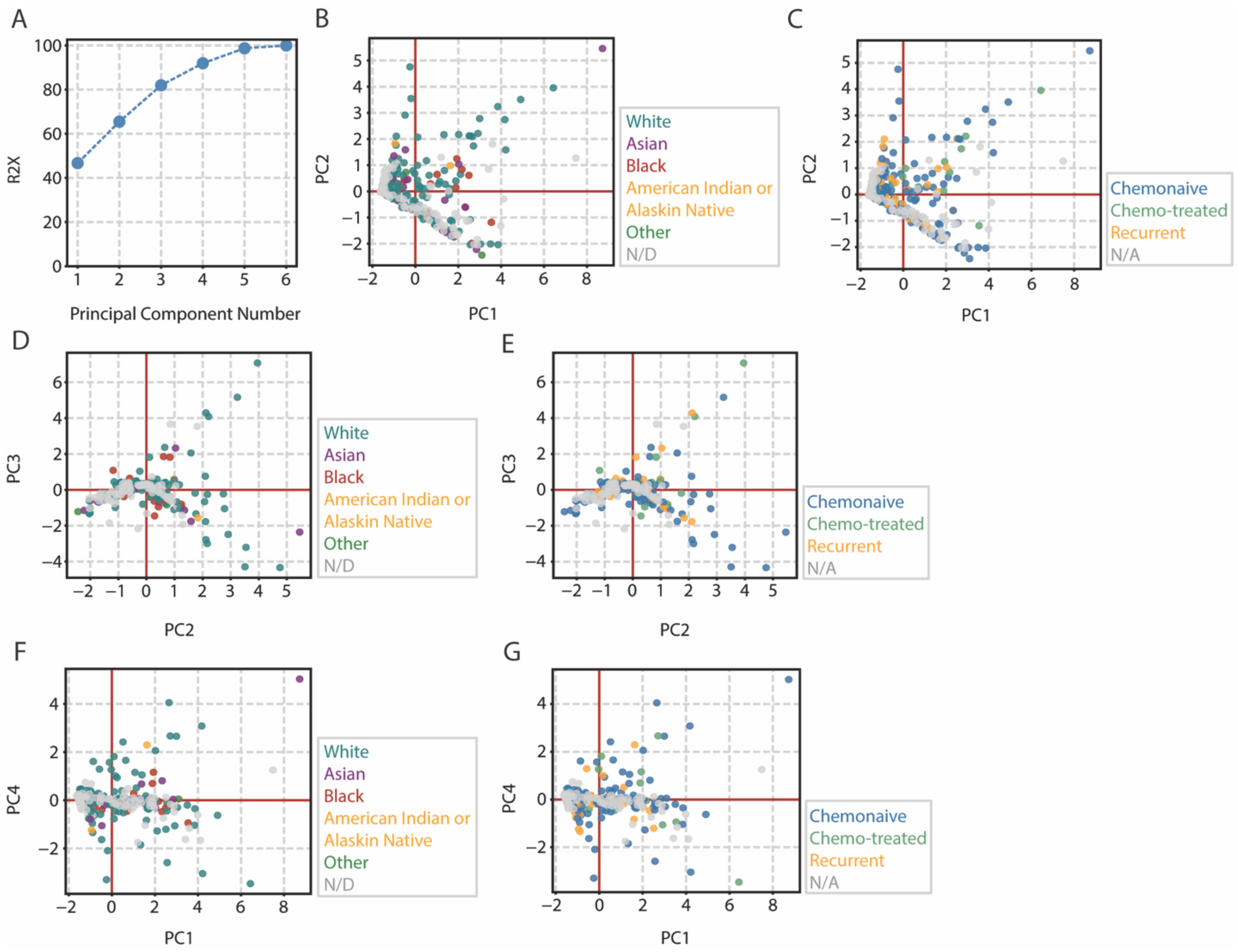


**Figure S3. PCA of patient demographics, related to Figure 3.**

A) Percent variance explained by number of components.

PCA scores plots for the PCs, colored by patient race (B, D, F) and disease treatment history (C, E, G) for each measurement (n=81).

N/D (not determined) refers to healthy samples for which there was no available data. N/A (not applicable) refers to healthy serum samples that did not undergo chemotherapy.

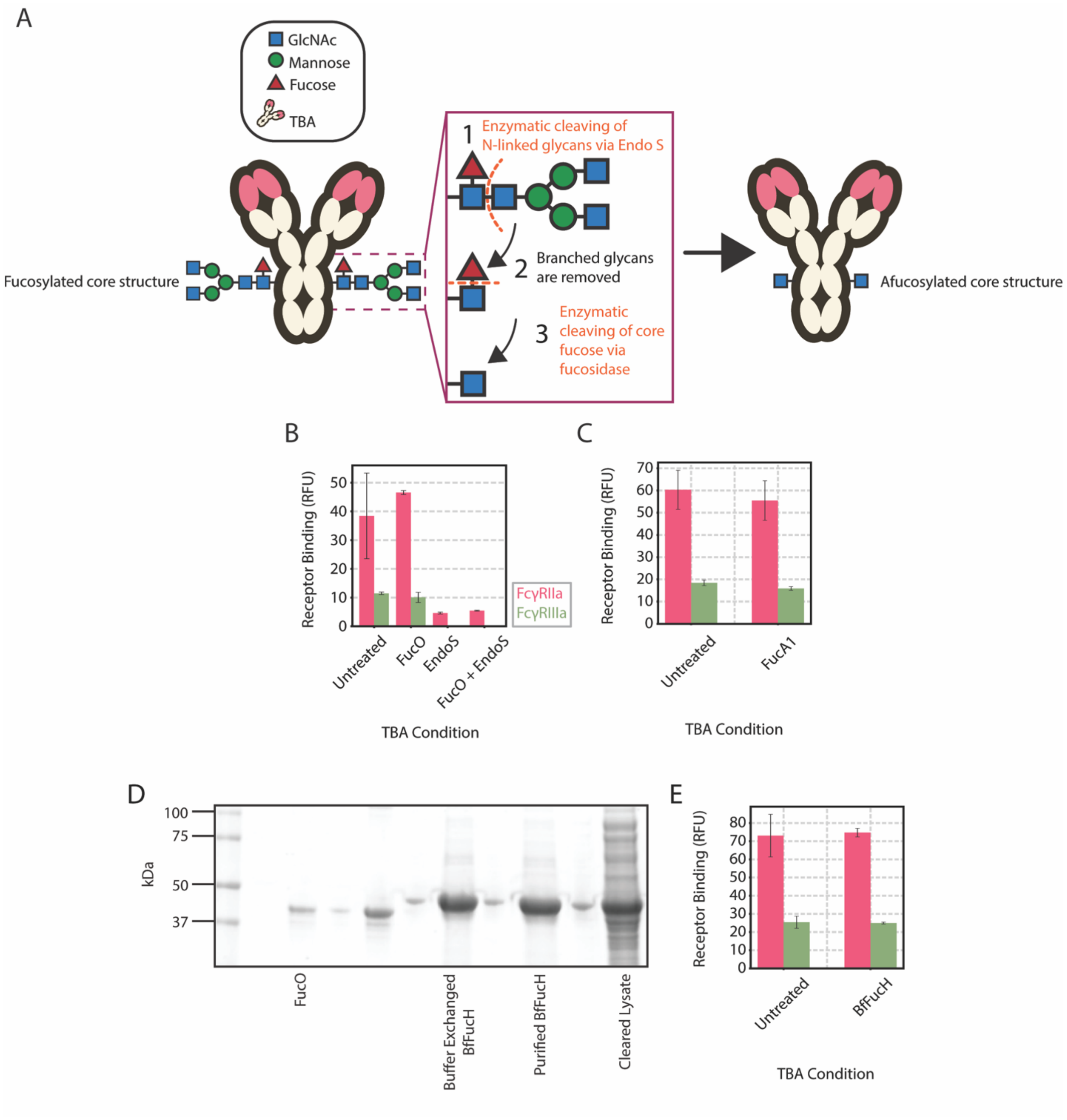


**Figure S4. Fucosidase treatment of TBAs, related to Figure 4.**

A) To evaluate enzyme activity, α1-2,4,6 Fucosidase O (FucO) was incubated with patient TBAs overnight, with or without simultaneous EndoS treatment, alongside a no-enzyme control.

B) FcγR binding of afucosylated TBAs, quantified by fluorescence (n=1).

C) FucA1 was incubated with patient TBAs for 7 days, alongside a no-enzyme control. FcγR binding was then quantified by fluorescence (n=1).

D) Coomassie stained gel showing raw lysate and purified BfFucH.

E) To evaluate enzyme activity, BfFucH was incubated with patient TBAs overnight, alongside an antibody-only control (n=1). FcγR binding was quantified by fluorescence.
